# A Structural Ensemble of a Tau-Microtubule Complex Reveals Regulatory Tau Phosphorylation and Acetylation Mechanisms

**DOI:** 10.1101/2020.11.10.376285

**Authors:** Z. Faidon Brotzakis, Philip R. Lindstedt, Ross Taylor, Gonçalo J. L. Bernardes, Michele Vendruscolo

## Abstract

Tau is a microtubule-associated protein that regulates the stability of microtubules. The affinity of tau for microtubules is modulated by post-translational modifications, and the dysregulation of these events has been associated with the aberrant aggregation of tau in Alzheimer's disease and related tauopathies. Here, we use the metainference cryo-electron microscopy approach to determine an ensemble of structures representing the structure and dynamics of a tau-microtubule complex comprising an extended microtubule-binding region of tau (residues 202-395). We thus identify the ground state of the complex and a series of excited states of lower populations. An analysis of the interactions in these states of structures reveals positions in the tau sequence that are important to determine the overall stability of the tau-microtubule complex. This analysis leads to the identification of positions where phosphorylation and acetylation events have destabilising effects, which we validate by using site-specific post-translationally modified tau variants obtained by chemical mutagenesis. Taken together, these results illustrate how the simultaneous determination of ground and excited states of macromolecular complexes reveals functional and regulatory mechanisms.

---

Microtubules are essential components of the cytoskeleton formed by the polymerisation of dimers of α-tubulin and β-tubulin, which are stabilised by a family of microtubule-associated proteins, of which tau is a member^1^. Tau has μM affinity to microtubules, primarily interacting through its microtubule binding domain (MBD) repeats (R1-R4), and through the flexible PGGG (P2) and proline-rich (R’) domains (**Fig. 1A**)^2,3^. The C-terminal region of β-tubulin also contributes to the stability of the tau-microtubule complex^4,5^. Recent advances in cryo-electron microscopy (cryo-EM) have recently led to the determination of the atomistic interactions of R1 and R2 with microtubules (**Fig. 1A**)^6^. Other regions of the tau-microtubule complex were determined at lower resolution (4.5-6.5 Å) due to conformational heterogeneity of the corresponding tau and microtubule regions, preventing a fully atomistic description of some of the stabilizing contacts, especially in the R3 and R4 repeat regions, the C-terminal region of β-tubulin, the flexible PGGG regions and the proline-rich domain.

**Figure 1.**
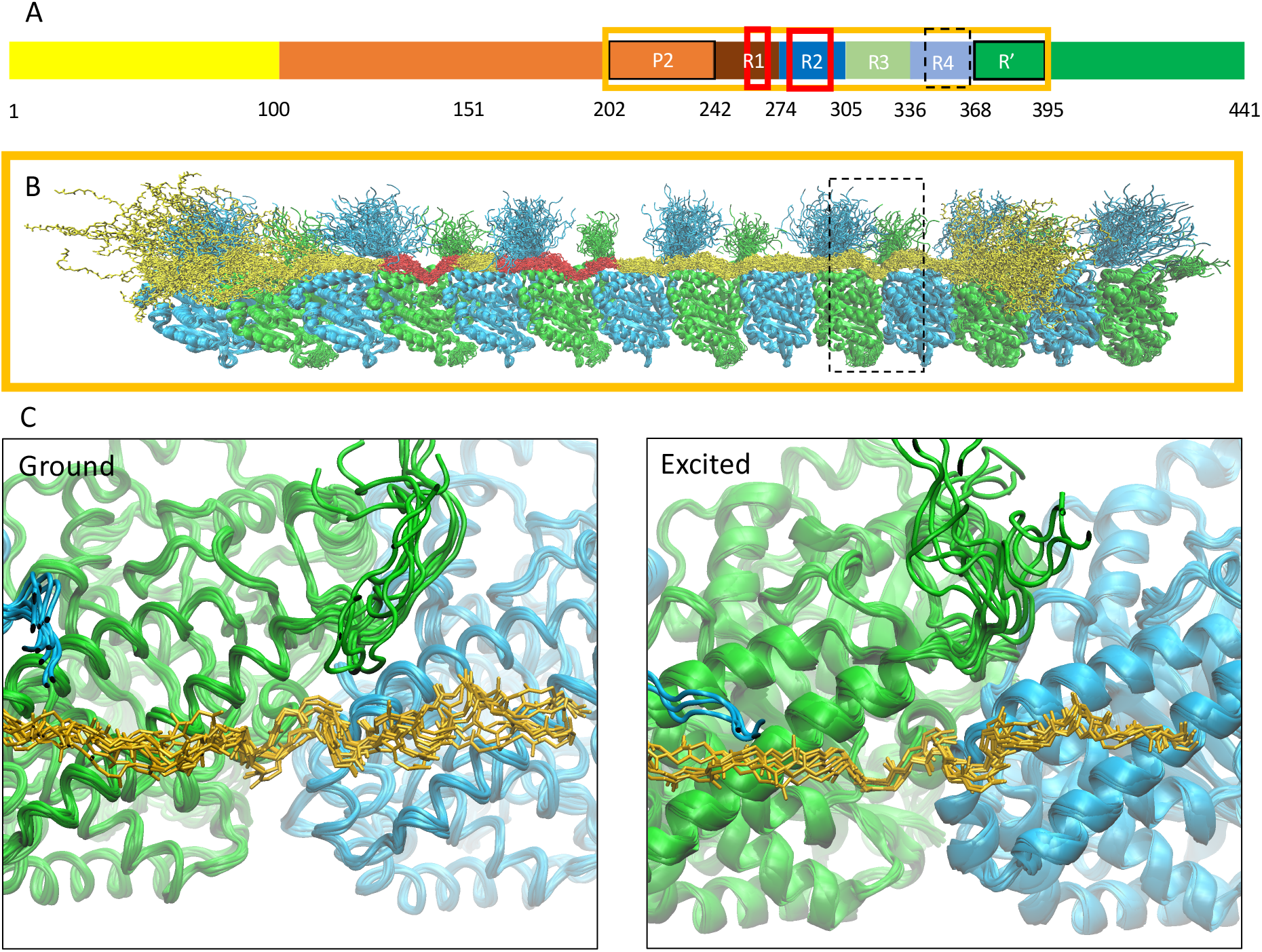
Simultaneous determination of the structure and dynamics of a tau-microtubule complex. **(A)** Illustration of the functional domains in the tau sequence, and sequence alignment of microtubule binding repeats (R1, R2, R3, R4) and flanking regions (P2, R’). The dashed and solid box sequence region is referred to the weak and strong region, respectively. The red solid bars below the R1 and R2 regions identify the regions whose structure was previously determined^6^ and the gold solid bar identifies the region whose structure is determined here (residues 202-395, comprising the regions P2-R1-R2-R3-R4-R’). **(B)** Atomic-resolution structural ensemble of an extended microtubule-binding region of tau (residues 202-395) in complex with a microtubule, as determined by EMMI in this study. α-tubulin, β-tubulin and tau coloured green, blue and gold, respectively. **(C)** Structural details of the R4 region in the black box in panel B. Since the structural ensemble contains multiple substates (**Fig. S2**) we show both the most populated state (ground state) and the second most populated state (excited state).

It is therefore important to characterise the dynamics of this macromolecular complex in order to better understand the structural basis for the stabilisation of microtubule by tau and for the regulatory role of tau phosphorylation and acetylation sites.

This goal can be achieved by using cryo-EM measurements to determine structural ensembles representing dynamics of the complex. A major challenge towards this goal is to disentangle in the measurements the effects of macromolecular dynamics from those of systematic and random errors. The recent development of the metainference method has enabled to overcome this challenge^7^. The goal of metainference is to accurately model a thermodynamic ensemble by optimally combining prior information of the system such as physico-chemical knowledge with experimental data, and by determining the level of noise in the experimental input data. Metainference has already been used successfully in a series of complex biological problems in combination with cryo-EM and other techniques^8,9^.

## Simultaneous determination of the structure and dynamics of a tau-microtubule complex

The tau-microtubule structural ensemble determined in this study (**Fig. 1B)** provides novel insight into the behaviour of the microtubule-binding region of tau (in particular P2, R3, R4, and R’) and of the tubulin C-terminal region. Notably, P2, R’ and the C-termini of tubulin are rather disordered, consistent with the weak densities that these regions have in the cryo-EM maps and NMR experiments^3,6^.

To illustrate the ability of the metainference cryo-EM (EMMI) method to disentangle the experimental error from the structural heterogeneity, we plot back-calculated EM maps generated only by the heterogeneity of the system - that is only by the structures of the EMMI ensemble - as a function of density as shown in **Fig. S1**. For high electron densities most of the microtubule structure is identifiable, indicating low levels of structural heterogeneity and a fairly rigid microtubule structure. However, as the electron density weakens (moving downwards in **Fig. S1**) structurally heterogeneous regions start to appear more clearly, such as R1-R4, P2 and R’ as well as the C-termini of tubulin monomers.

The dynamics of the tau-microtubule complex results in the population of different states. Our results indicate that the complex populates a ground state, as well as several excited states at lower populations (**Fig. S2**), which we obtained by performing a clustering analysis of the structural ensemble (**Fig. 1B**). We compare the structures of the ground state and an excited state for the otherwise undetermined R4 repeat in **Fig. 1C**.

## Stabilising regions in the tau-microtubule complex

The tau-microtubule complex exhibits highly dynamical regions characterised by large conformational fluctuations (**Fig. 2A**). In the complex, tau has a rigid microtubule binding domain region and two flexible flanking regions (P2 and R’), with the P2 region exhibiting greater structural heterogeneity (4.33 ± 0.05 nm) than R’ (3.20 ± 0.04 nm). We also note that the R1 region exhibits slightly, but significantly, greater structural heterogeneity (0.56 ± 0.02 nm) than the R2 (0.43 ± 0.02 nm), R3 (0.42 ± 0.01 nm) and R4 (0.50 ± 0.02 nm) regions. Such evidence suggests the possibility of an asymmetric unzipping mechanism, where tau preferentially unbinds from the microtubules from the P2 region rather than from the R’ region. To further shed light on this mechanism, we discuss the tau-microtubule interaction energies quantified by the contacts each tau residue forms with the microtubule (**Fig. 2B**). We thus identify weak and strong interacting regions of tau with microtubules, indicated by dashed and solid arrows. For each repeat, we associate weakly (R1w, R2w, R3w, R4w) and strongly (R1s, R2s, R3s, R4s) interacting regions, which correspond to residue sequences indicated by the dashed and solid boxes in **Fig. S3A** respectively. The strong interacting area of the repeats includes the hallmark SKI(C)GS motif, known to contribute to tau-microtubule stability and be associated with AD related phosphorylation sites^10,11^. Notably, the weak interacting region also shows some sequence conservation motif across the repeats with the PHF6 aggregation prone VQIIN(VY)K hallmark motif^12^ present in the R2 and R3 regions. As described below, lysine residues of the weak regions play a critical role in stabilizing the interaction with microtubules. Residues 230-240 (P2w) and 370-379 (R’w) form weak interactions with the microtubule, with serine residues in the P2w region and lysine ones in the R’w region contributing to the tau-microtubule interaction.

**Figure 2.**
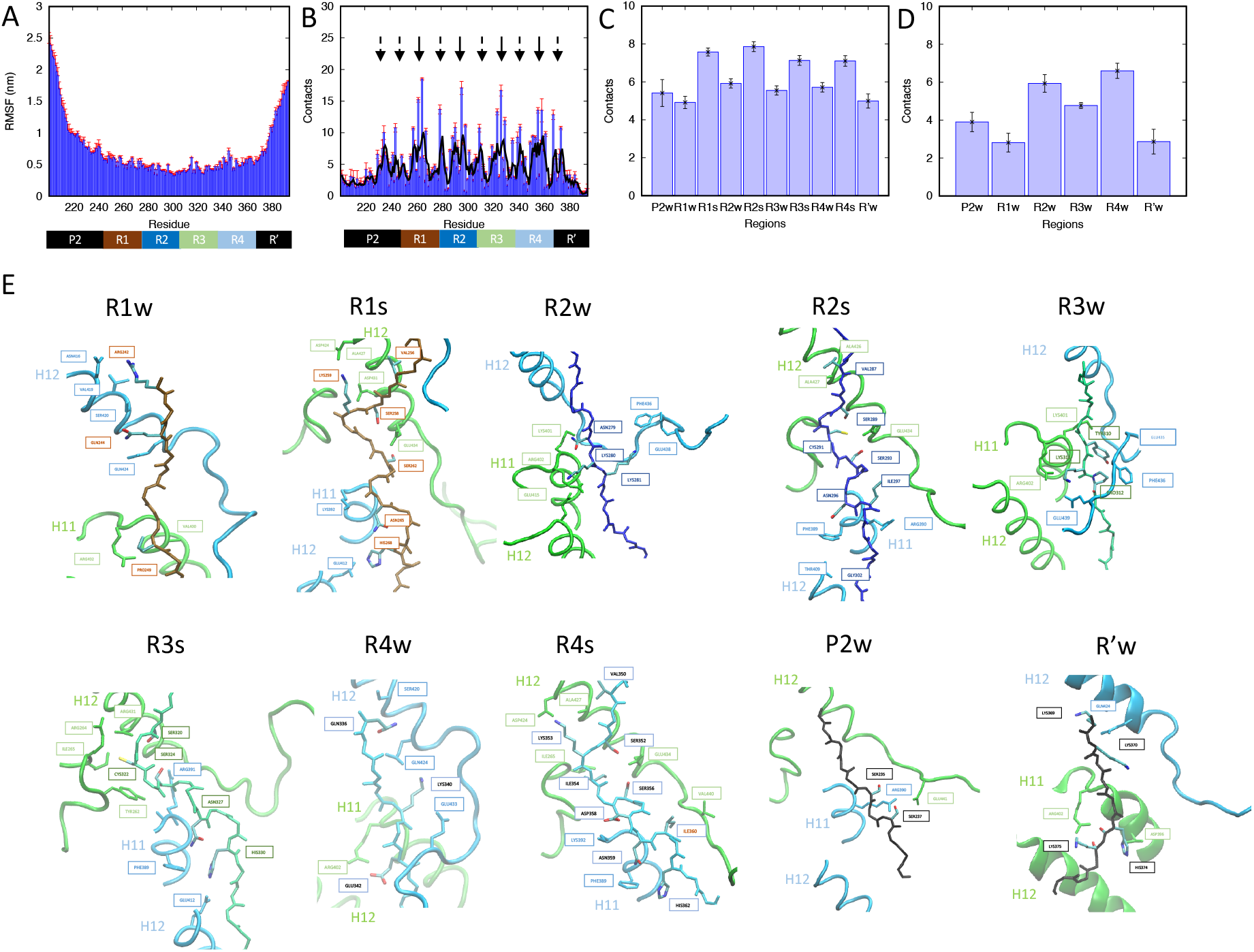
Stabilising regions in the tau-microtubule complex. **(A)** All-atom root mean square fluctuations along the tau sequence. **(B)** Number of contacts between each tau residue and the microtubule. The black curve indicates a running average with a 3-residue window. **(C)** Average number of tau-microtubule contacts per tau region and per residue (normalized by the number of respective regions in the region). **(D)** Average number of contacts between tau and α-tubulin β-tubulin C-terminal per tau weakly interacting region and per residue (normalized by the number of respective regions in the weak region). **(E**) Weakly and strongly interacting regions of the most populated structure as obtained from the clustering analysis.

Each residue of the weak or strong regions forms on average about 5-6 or 7.5-8 contacts with microtubules respectively (**Fig. 2C**), with a repetitive pattern of weak interactions followed by strong interactions in the microtubule binding domain region. Notably, the weak interaction region of R1 (R1w) forms slightly fewer contacts with the microtubules than other weak regions do. These observations are attributed to the fewer interactions R1w region makes with the C-terminal of β-tubulin as opposed to the R2w, R3w, R4w regions (**Fig. 2D**) and can explain the higher flexibility of the R1 region with respect to rest repeat regions. Evidently, the β-tubulin C-termini stabilize the tau-microtubule complex by forming interactions with the weak regions and this could explain the stabilizing role of C-termini in tau-microtubule complex^4,5^.

Overall, the higher flexibility of the P2 region with respect to the R’ region, and of the R1 region with respect to the R2-R4 regions, the fewer contacts that the R1w region makes with microtubules with respect to the R2w, R3w, and R4w regions to microtubules, may suggest an asymmetric unzipping mechanism arising preferentially from the P2 region rather than the R’ region. In an unzipping from R4 region, tau would first have to break the strong interactions in the R4 region, while in an unzipping from the R1 region, tau would first have to break the weak interactions in the R1 region.

## Interatomic interactions stabilising the tau-microtubule complex

Tau interacts in a similar manner with tubulin along the strongly interacting regions (R1s, R2s, R3s, R4s), in particular with α-helix H12 of α-tubulin and α-helices H11 and H12 of β-tubulin (**Fig. 2E**). The weakly interaction regions (R1w, R2w, R3w, R4w, R’w) also interact in a similar manner, namely with α-helix H12 of β-tubulin, and α-helices H11 and H12 of β-tubulin. The only exception to this interaction symmetry is the P2w region, which interacts with parts of tubulin that a strong interacting region would have, i.e with α-helix H12 of α-tubulin as well as with α-helices H11 and H12 of β-tubulin. This is obligatory action for tau since the entire KIGS…PGGG sequence domain existing in the microtubule-binding repeats is missing in the P2 region (**Fig. S3A**), and therefore residues in the weakly interacting regions replace such interactions. This mismatch apparently leads to weaker interactions of tau P2 region with microtubules.

The strongly interacting region is characterized by a conserved sequence identity (**Fig. S3A**), as well as interaction between amino acids of tau and tubulin, from repeat to repeat. V256, V287, V316 and V350 make hydrophobic interactions with A427 in α-tubulin. The residues of the SKI(C)GS motif interact as follows with microtubules: residues S258, S289, S352 and S320 form contacts with residues E434, K259, K290, K321, K353 in α-tubulin, residues D424, D431 and S262, S293, S324, S356 with residue E434 in α-tubulin. Finally, the sequence preserved N265, N296, N327, N359 and H268, H330, H362 interact with residues K392, F398 and E412 in β-tubulin, respectively. The experimentally identified^10,11,13,14^ important contribution of S262, S324, S356, K259, K290, K321, K353 in the stability of the tau-microtubule complex concurs with our findings.

In the weak interaction region R1w, residue R242 of tau is in contact with residues N416 and S420 of α-tubulin, and residue Q244 of tau with residue S420 in α-tubulin. In the weakly interacting region R2w, residues N279, K280 and K281 of tau form salt-bridge interactions with residues K401, R402, E414 of α-tubulin and the C-terminal residue E438 of β-tubulin. In the weakly interacting region R3w, residues Y310, K311 and P3121 of tau form a combination of salt-bridge or apolar interactions with C-terminal residues F436, E439 of β-tubulin and residues R401, K402 of α-tubulin. Notably, residues K280, K281, Y310, K311 are part of the VQIIN(VY)K sequence preserved motif with residues K280, K281 known to contribute to the stability of the tau-microtubule complex while the entire motif is an aggregation prone motif^12,14^. The R4w region comprises notable interactions between residues Q336, K340 and E342 of tau with residues S420, N424, E433 of β-tubulin and R402 of α-tubulin. The R’w region forms notable interactions between residues K369, K370, K375 and H374 of tau with residues N424 of β-tubulin and R402 and D396 of α-tubulin. Finally, in the P2w region, residues S235, S237 of tau interact with residues E441 of α-tubulin and R390 of β-tubulin.

These results are in agreement with previously determined structures of the R1 and R2 regions^6^ (**Fig. S3B,C**) within an RMSD (backbone and sidechains) of 0.27 ± 0.15 Å for the R1 region and 0.3 ± 0.2 Å for the R2 region from different structures of the most populated cluster of our EMMI ensemble.

## Identification and validation of phosphorylation sites altering the stability of the tau-microtubule complex

The phosphorylation of tau can lead to a loss of stabilizing tau-microtubule interactions. Having access to the structure and dynamics of the tau-microtubule complex we can make a connection between the strength of the tau-microtubule interactions and the propensity of residues to be phosphorylated. Such a connection can not only verify existing phosphorylation sites altering the tau-microtubule stability, but also predict new ones. We analysed the average number of contacts and the conformational heterogeneity of serine and tyrosine residues (**Fig. 3A**). According to this metric, residues that contribute most to the stability are those that form high number of contacts, and residues that have high conformational heterogeneity have a higher chance to become accessible to kinases and be post-translationally modified at the tau-microtubule interface. This approach is validated by identifying known tau-microtubule stability-altering phosphorylation sites in AD brains (**Fig. 3A**, orange sites), including S258, S262, S324, S356^10,13,15,16^, since they form many contacts with microtubules and have a moderate flexibility. In addition, this analysis predicts known AD brain phosphorylated sites with unknown tau-microtubule stability role, including S237 (P2w) and S289 (R2s) (**Fig. 3B**, orange sites)^11^.

**Figure 3.**
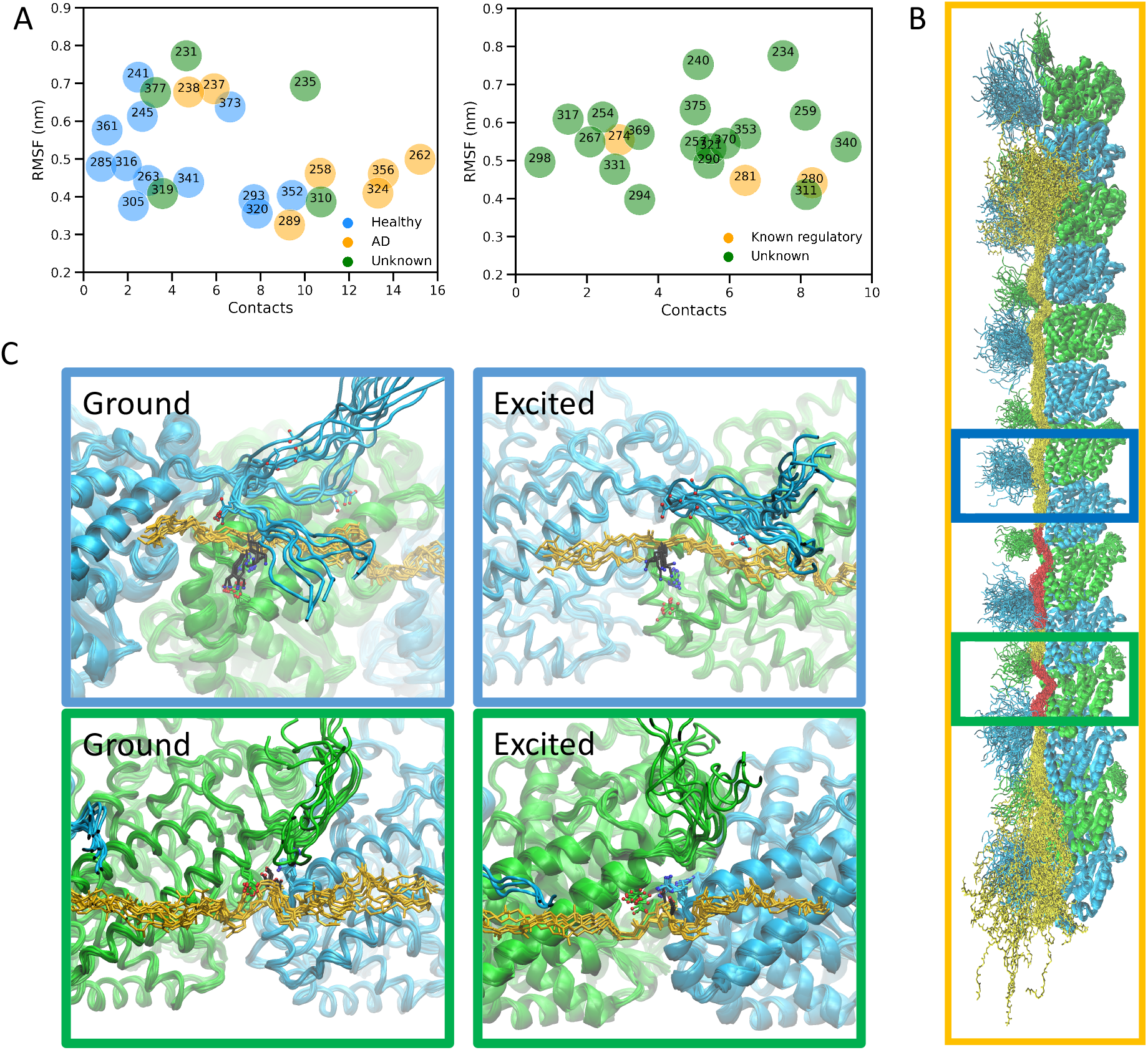
Identification of phosphorylation and acetylation sites altering the stability of the tau-microtubule complex. **(A)** By considering the various contributions to the stability of the tau-microtubule complex (number of contacts of a residue, x-axis) and its accessibility to kinases (structural heterogeneity, RMSF, y-axis), we can identify the residues in the upper right region of the plot as those that are most important for stability and have the higher propensity to be phosphorylated or acetylated. Our results show that most residues known to be phosphorylated in AD (in orange) are of this type. Residues phosphorylated in healthy brains (in blue) tend instead to have fewer contacts and therefore to have a weaker effect on stability. From this analysis, we identify S262 as potentially having a significant effect on the stability. Phosphorylation data are obtained from Ref.^11^. Similarly, we can identify the residues that are most important for stability and have the higher propensity to be acetylated. Our results show that two of the residues with regulatory function, known to be acetylated in AD (K280 and K281, in orange) are of this type. Among the residues whose acetylation state is not known (in green), we identify K311 as potentially having a significant effect on the stability. Acetylated data are obtained from^29,30^. **(B)** Positions of the regions of residues S262 (green) and K311 (blue), which we identified for post-translational modifications. **(C)** Details of the structural ensembles showing the interactions of tau S262 and tubulin E434, and of tau K311 with tubulin R402 and E436. The comparison between the structures of the ground states and the excited states illustrates the availability in the excited state of the sites for post-translational modifications.

Next, we analysed in more detail the phosphorylation at position S262 (**Fig. 3B**, **green box**). The excited state shows that this position becomes more exposed and hence possibly more accessible to kinases for phosphorylation (**Fig. 3C**, **green box**). We validate these calculations by a microtubule-binding assay (**Fig. 4A,B**), which shows that phosphorylation of S262 reduces the microtubule polymerization rate. In order to obtain site-specific phosphorylation at S262, we use an established chemical mutagenesis approach that has been used to accurately mimic phosphorylation on histones and kinases^17,18^. This approach relies on the installation of the amino acid dehydroalanine (Dha) at the site of interest and then a subsequent Michael-addition by thiophosphate to install a highly accurate phosphorylation mimic (**Fig. S4**). Importantly, we also identified sites with unknown phosphorylation state in AD brains and unknown tau-microtubule stability role such as S235 (P2w) and Y310 (R2w) (**Fig. 3A**, green sites)^11^. Finally, we observe that phosphorylated residues in healthy brains did not contribute much to the stability of the tau-microtubule complex since mostly form few interactions with microtubules (**Fig. 3A**, blue sites).

**Figure 4.**
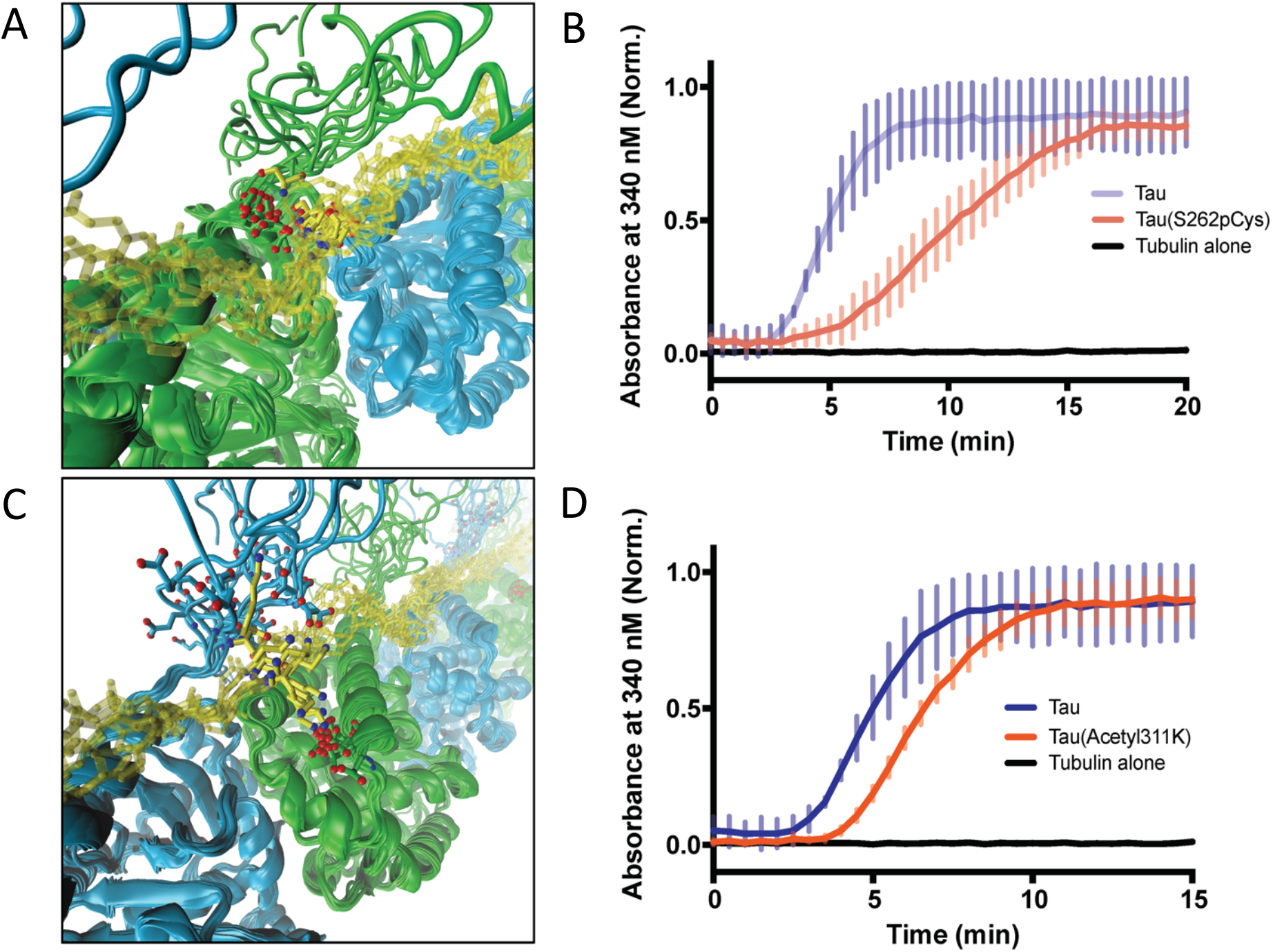
Validation of phosphorylation and acetylation sites altering the stability of the tau-microtubule complex. **(A)** Conformational heterogeneity of the interaction of tau S262 with tubulin E434. **(B)** Validation of the effect of phosphorylation of S262 on tubulin polymerisation. **(C)** Conformational heterogeneity of the interaction of tau K311 with tubulin R402 and E436. **(D)** Validation of the effect of acetylation of K311 on tubulin polymerisation.

## Identification and validation of acetylation sites altering the stability of the tau-microtubule complex

Tau acetylation in relation to tau-microtubule stability dysregulation is less studied than phosphorylation. In a similar manner as in the previous paragraph, we select tau-microtubule stability altering upon acetylation-prone residues i.e. forming many contacts and exhibit moderate to high flexibility (**Fig. 3A**). This analysis identifies K280, K281 and K274 known sites which upon acetylation significantly dysregulate the complex stability (**Fig. 3A**, orange sites), with the first two being related to AD pathology^14^. The analysis also predicts K234 (P2w), K240 (P2w), K259 (R1s), 290 (R2s), 311 (R3w), 321 (R3s), 340 (R4w), 353 (R4s), 370 (R’w), 375 (R’w) as affinity-regulating acetylation sites (**Fig. 3A**, green sites). K259, K290, K321, K353 belong to the KXGS domain, while stronger contacts such as 280 (R2w), 31 (R3w), 340 (R4w) are repeated lysine residues on the weak interaction region of tau VQIIN(VY)K. Next, we analysed in more detail the phosphorylation at position K311 (**Fig. 3B**, **blue box**). We then validated these predictions by a microtubule-binding assay (**Fig. 4C,D**), which shows that acetylation of K311 reduces the microtubule polymerization rate. In order to obtain site-specific acetylation at K311 we used the same chemical mutagenesis approach as taken with the phosphorylation mimetics except that the Michael-addition to Dha at position 311 was carried out by *N*-acetylcysteamine (**Fig. S5**).

In conclusion, we have reported a structural ensemble of a tau-microtubule complex. The results that we have presented reveal how the conformational fluctuations in the complex lead to the population of excited states. We then identify post-translational modification sites that take part in the regulation of the binding of tau to microtubules, and validate their effects on microtubule polymerisation through a site-specific chemical mutagenesis approach. These results thus provide a mechanistic understanding of one of the molecular processes most closely linked to the progression of Alzheimer’s disease^10,11,14,19^.

## Acknowledgements

The authors would like to acknowledge Dr. Massimiliano Bonomi, Dr. Liz Kellogg and Dr. Eva Nogales for discussions, and ARCHER supercomputer system for making computer time available. Z.F.B. would like to acknowledge the Federation of European Biochemical Societies (FEBS) for financial support (LTF). G. J. L. B. is a Royal Society University Research Fellow (URF/R/180019).

## Methods

### The EMMI method

Metainference^7^ has been extended to work with cryo-EM data in the EMMI method^8^. In EMMI, the cryo-EM data voxel map is represented as a Gaussian Mixture Model (GMM) *ϕ*_*D*_ with *N*_*D*_ components

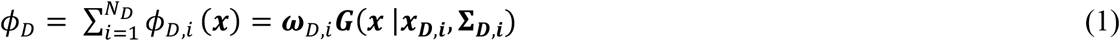

where ***ω***_*D,i*_ is the weight of the ith component of the data GMM and ***G*** is a normalized Gaussian function centered in ***x***_*D,i*_ with a corresponding covariance matrix **Σ**_***D,i***_. EMMI quantifies the deviation between the experimental data GMM and the molecular dynamics generated models GMM by using the following overlap function

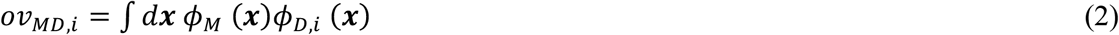

where *ϕ*_*M*_ (***x***) corresponds to the model GMM. Since EMMI deals with the heterogeneity of the system by simulating many replicas of it, the overlap between model GMM and data GMM is estimated over the ensemble of replicas to an average overlap 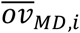. Finally, since cryo-EM maps usually contain big amount of density particles data, EMMI samples the error in the data *a posteriori*, thus simplifying the total energy function to

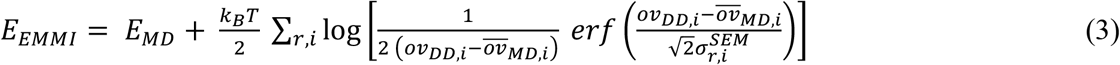

where the first energy term corresponds to the force field and the second energy term provides with an energy penalty depending on the agreement of the data with the molecular dynamics generated models.

### Structural ensemble calculations setup

We start by building a microtubule segment comprising seven α-tubulin/β-tubulin dimers, obtained from the cryo-EM characterized structure (PDB:6CVJ). We continue with constructing an initial structure for tau (residues 202-395), where tau-microtubule binding domain regions R1-R2 are based on PDBs 6CVJ and 6CVN. The interactions of R1-R2 with the microtubule are used as a template to build the initial structure of tau region R3-R4 while we impose the R3-R4 structure should fit in the full-length tau cryo-EM map density (EMD-7522). We achieve so by first fitting a polyalanine model in Rosetta package^20^ and then adding side-chains. Residues 202-241 (P2 region) and 369-395 (R’ region) are built with a random initial structure as these regions are flexible as indicated by the absence of strong density in the tau-microtubule full-length cryo-EM map.

### Molecular dynamics equilibration

We continue with setting a 9.7 × 11.4 × 63.6 nm simulation box, solvating with 202,493 water molecules and neutralizing it by adding ions. We use the AMBER99SB-ILDN^21^ and TIP3P^22^ protein and water forcefield respectively. We carry on with an energy minimization step followed by a short NPT simulation, followed by a short NVT simulation.

All bonds are constrained with the LINCS algorithm^23^. A cut-off value of 1 nm is used for the Lennard-Jones interactions. The Particle Mesh Ewald method is used to calculate the electrostatic interactions with a Fourier spacing 1.2 nm and 1 nm cut-off for the short-range electrostatic interactions. The pair lists are updated every 10 fs with a cut-off of 1 nm and the timestep was 2 fs^22^. A leap-frog algorithm for integrating Newton’s equations of motion is used, the velocity-rescale thermostat^24^ with a coupling time constant of 0.2 ps, and Parrinello-Rahman barostat^25^ with a coupling time constant of 1.0 ps is used for performing an NPT simulation. In the NPT, Cα are position restrained with a constant 200 kJ/mol nm^2^, temperature is set to 310 K, pressure to 1 atm and the simulation duration to 500 ps. In the NVT simulation position restraints are lifted, the simulation duration is 2 ns and temperature is set to 310 K without pressure coupling.

### EMMI simulations

First, we expressed the experimental voxel map data as a data GMM containing 47,157 Gaussians in total showing 0.95 correlation to the original voxel experimental map. We carry on by extracting 32 configurations from the previous NVT step and initiate two individual EMMI simulations, each consisting of 32 replicas and aggregate runtime 400 ns using PLUMED.2.6.0-dev^26^. EMMI simulations are performed in the NVT ensemble using the same MD parameters as in the equilibration step. Configurations were saved every 5 ps for postprocessing. The cryo-EM restraint is calculated every 2 MD steps, using neighbour lists to compute the overlaps between model and data GMMs, with cutoff equal to 0.01 and update frequency of 100 steps. In each EMMI simulation we toss the first 2 ns, divide the rest in two segment and perform cluster analysis using the gromos method on side-chain and backbone atoms and a 0.3 nm cut-off. For convergence purposes, in each EMMI simulation, we calculate cluster population averages and errors based on these two simulation segments. For molecular visualizations we use VMD^27^ and Chimera^28^.

### Tau expression and purification

2N4R tau lacking the endogenous cysteine residues (C291S & C322S) and the relevant cysteine mutants (S262C and K311C), created by standard site-directed mutagenesis, were expressed from a pet29b vector in BL21 Gold (DE3) cells (Agilent Technologies). Cultures were grown to an OD600 of 0.6 then induced with 0.4 mM IPTG and left to express at 18 °C overnight. Cells were harvested by centrifugation, resuspended in 50 mM MES (pH 6.5), 5 mM DTT, 0.1 mM PMSF and lysed via sonication (1 min 30 s; 5 s on, 10 s off; 40% amplitude) on ice. The lysed mixture was centrifuged, and tau was isolated via cation exchange using a Hitrap CaptoS column (GE Healthcare LifeSciences, Little Chalfont, U.K). Fractions containing tau as determined by gel electrophoresis were pooled and precipitated by the addition of 20% (w/v) ammonium sulphate on ice overnight. The precipitated protein was pelleted by centrifugation and then resuspended in SSPE buffer containing 5 mM DTT. Pure tau was finally isolated via size exclusion chromatography using a Superdex 200 Increase 10/300 GL column (GE Healthcare LifeSciences, Little Chalfont, U.K.) equilibrated with the aforementioned SSPE buffer, only the purest fractions as assessed by gel were kept for experiments.

### Dha formation

The tau cysteine mutants were buffer exchanged into 20 mM NaPi buffer (pH 8) via 7k MWCO Zeba spin desalting columns (Thermo Fisher). 200 μL of 50 μM protein aliquots were reacted with 50 molar equivalents of methyl 2,5-dibromopentanoate (Sigma Aldrich) for 12 hours at 37 °C and shaking at 500 rpm. Excess methyl 2,5-dibromopentanoate was removed by passing the reactions through 7k MWCO Zeba spin desalting columns and then conversion to Dha was verified via LC-MS

### Final chemical mutagenesis

For the creation of the phosphorylation mimetics, 100 μL aliquots of 50 μM of S262Dha in 20 mM NaPi (pH 8) were reacted batchwise (5 min intervals) with 30,000 molar equivalents of sodium thiophosphate (pH 8.0, 690 mg/mL suspension, 5×5000 equivalents). The mixtures were left to react for 8 hrs at 37 °C and shaking at 500 rpm, excess sodium thiosulphate was removed via two 7k MWCO Zeba spin desalting columns and reaction completion was verified via LC-MS. For the creation of the acetylated mimetic at K311, 100 μL aliquots of 50 μM K311Dha in in 20 mM NaPi (pH 8) was reacted with 1000 molar equivalents of *N*-acetylcysteamine (Sigma Aldrich) for 12 hours at 37 °C and shaking at 500 rpm. Excess *N*-acetylcysteamine was removed in the same manner as sodium thiophosphate and completion was again verified by LC-MS.

### Protein LC-MS

Protein LC–MS was performed on a Xevo G2-S TOF mass spectrometer coupled to an Acquity UPLC system using an Acquity UPLC BEH300 C4 column (1.7 μm, 2.1 mm × 50 mm). Water with 0.1% formic acid (solvent A) and 95% MeCN and 5% water with 0.1% formic acid (solvent B) were used as the mobile phase at a flow rate of 0.2 mL/min. The gradient was programmed as follows: 95% A for 0.93 min, then a gradient to 100% B over 4.28 min, then 100% B for 1.04 minutes, then a gradient to 95% A over 1.04 min. The electrospray source was operated with a capillary voltage of 2.0 kV and a cone voltage of 40 V. Nitrogen was used as the desolvation gas at a total flow of 850 L/h. Total mass spectra were reconstructed from the ion series using the MaxEnt algorithm preinstalled on MassLynx software (v4.1 from Waters) according to the manufacturer’s instructions.

### Microtubule polymerisation assay

The microtubule polymerisations were all performed with reagents from the kit purchased from Cytoskeleton, Inc. (Cat #BK006P) as per manufacturer’s instructions. Tubulin as added to the reaction mixture at a concentration of 3 mg/ mL and tau and the various chemical mutants were assayed at 15 μM for their ability to polymerize the reaction, which was monitored by OD at 340 nM using a CLARIOstar Plus plate reader (BMG Labtech).

---

**Figure S1.**
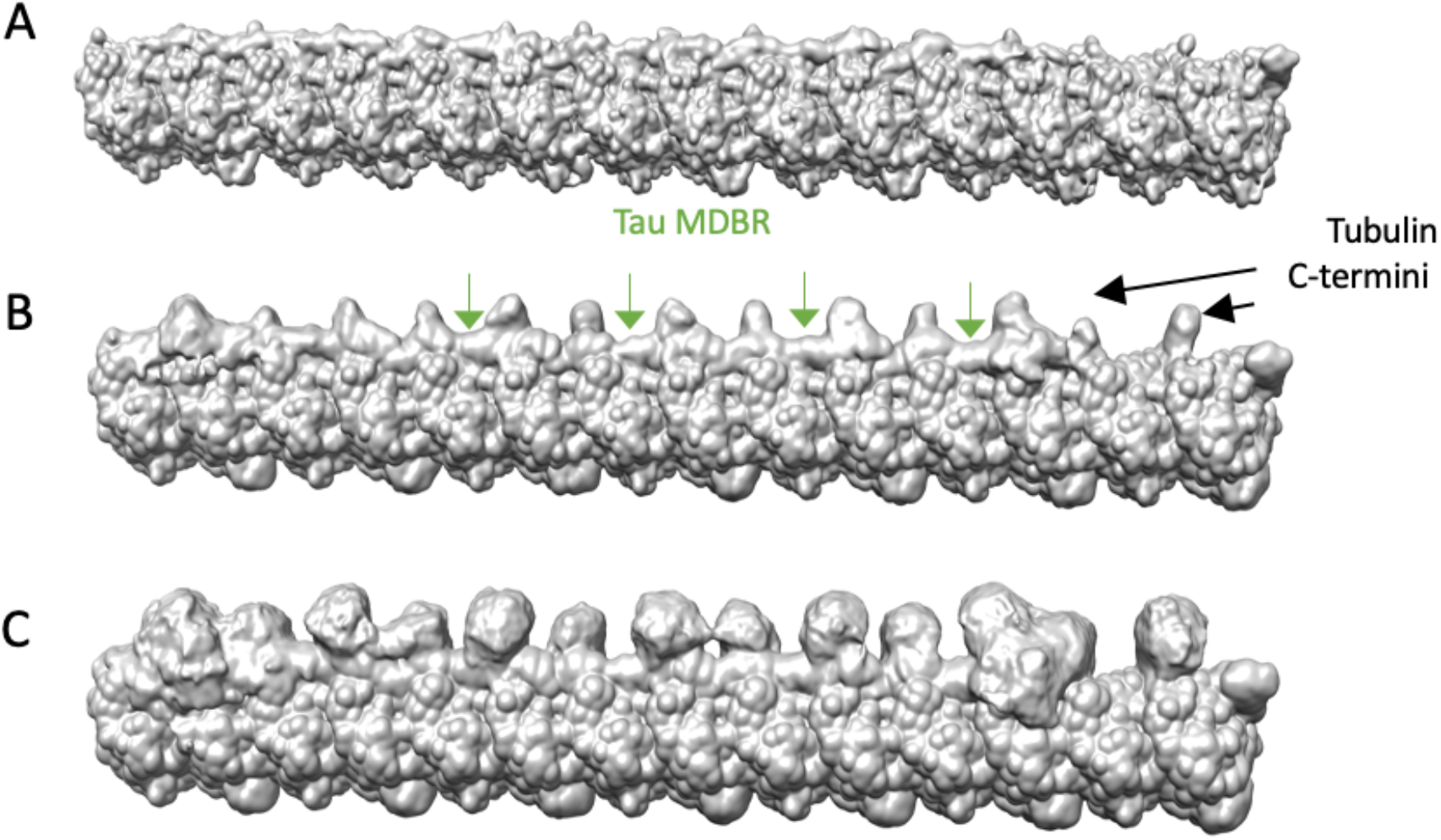
Back-calculated density maps from the EMMI ensemble as a function of decreasing electron density: (A) threshold 1, (B) threshold 0.5, and (C) threshold 0.05.

**Figure S2.**
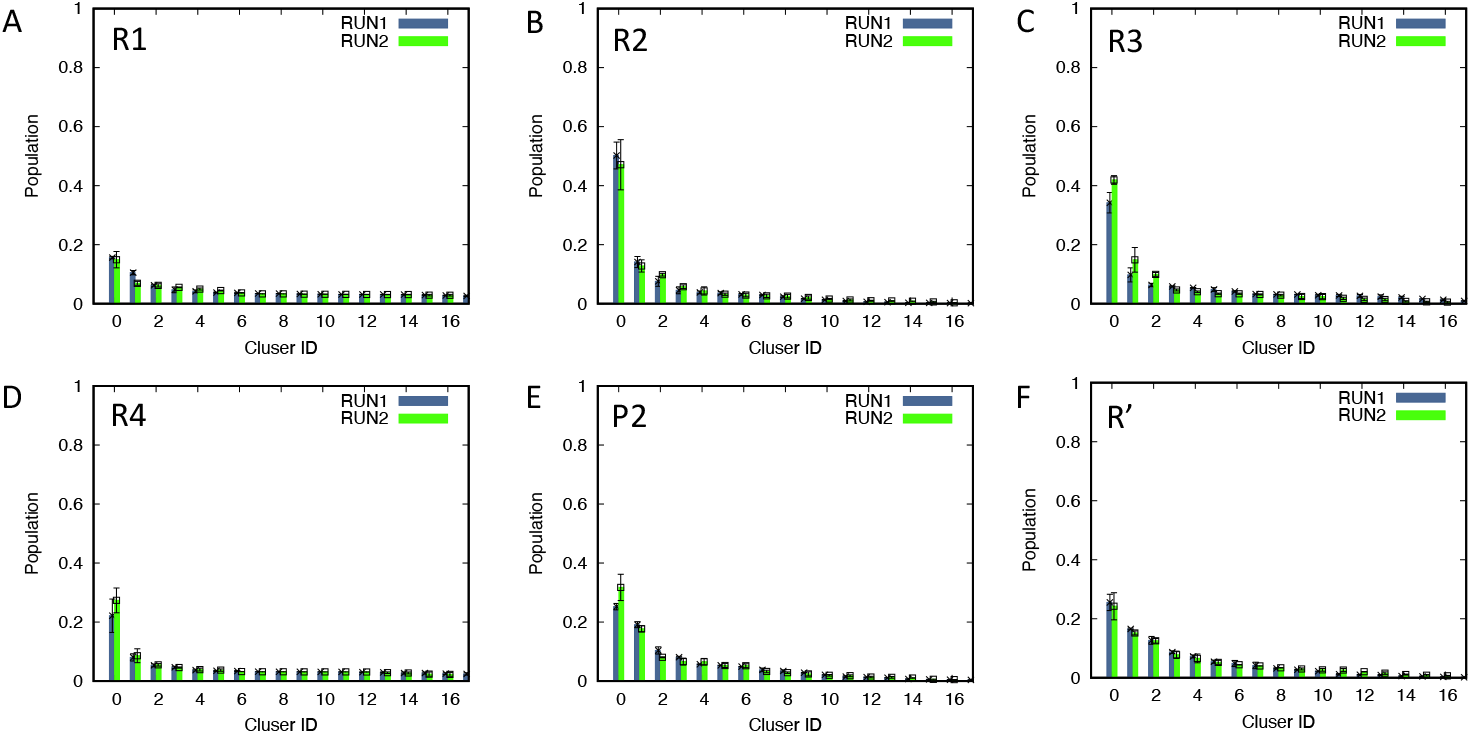
Convergence analysis of the EMMI structural ensemble of the R1 (A), R2 (B), R3 (C), R4 (D), P2 (E) and R’ (F) regions of tau.

**Figure S3.**
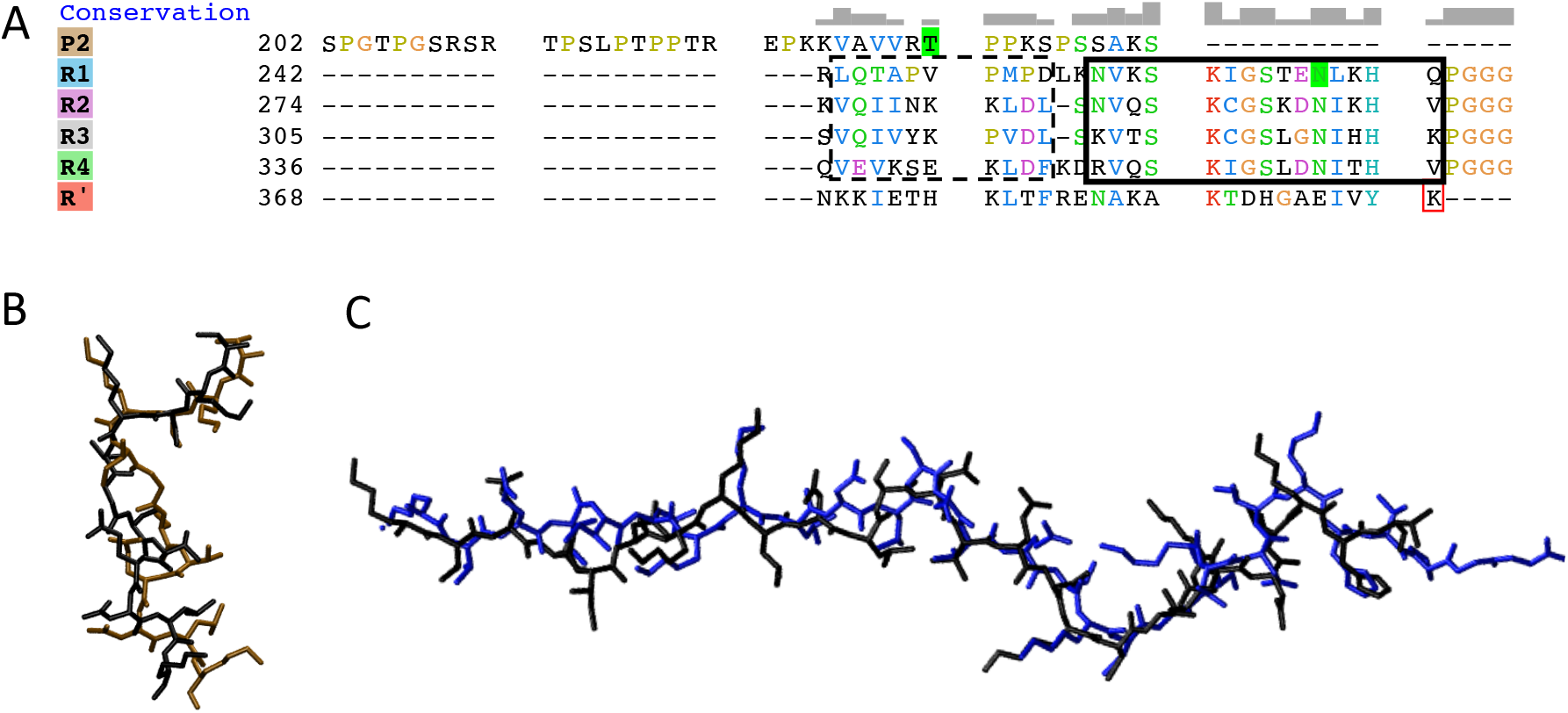
(A) Sequence alignment of the tau regions P2, R1, R2, R3, R4 and R’. With dashed and solid boxes are depicted the weak and strong interacting region of tau with the microtubules. (B) Comparison between the structures of the R1 region obtained in this work (in brown) and previously reported^6^ (black). (C) Comparison of the structures of the R2 region obtained in this work (in blue) and previously reported^6^ (black).

**Figure S4.**
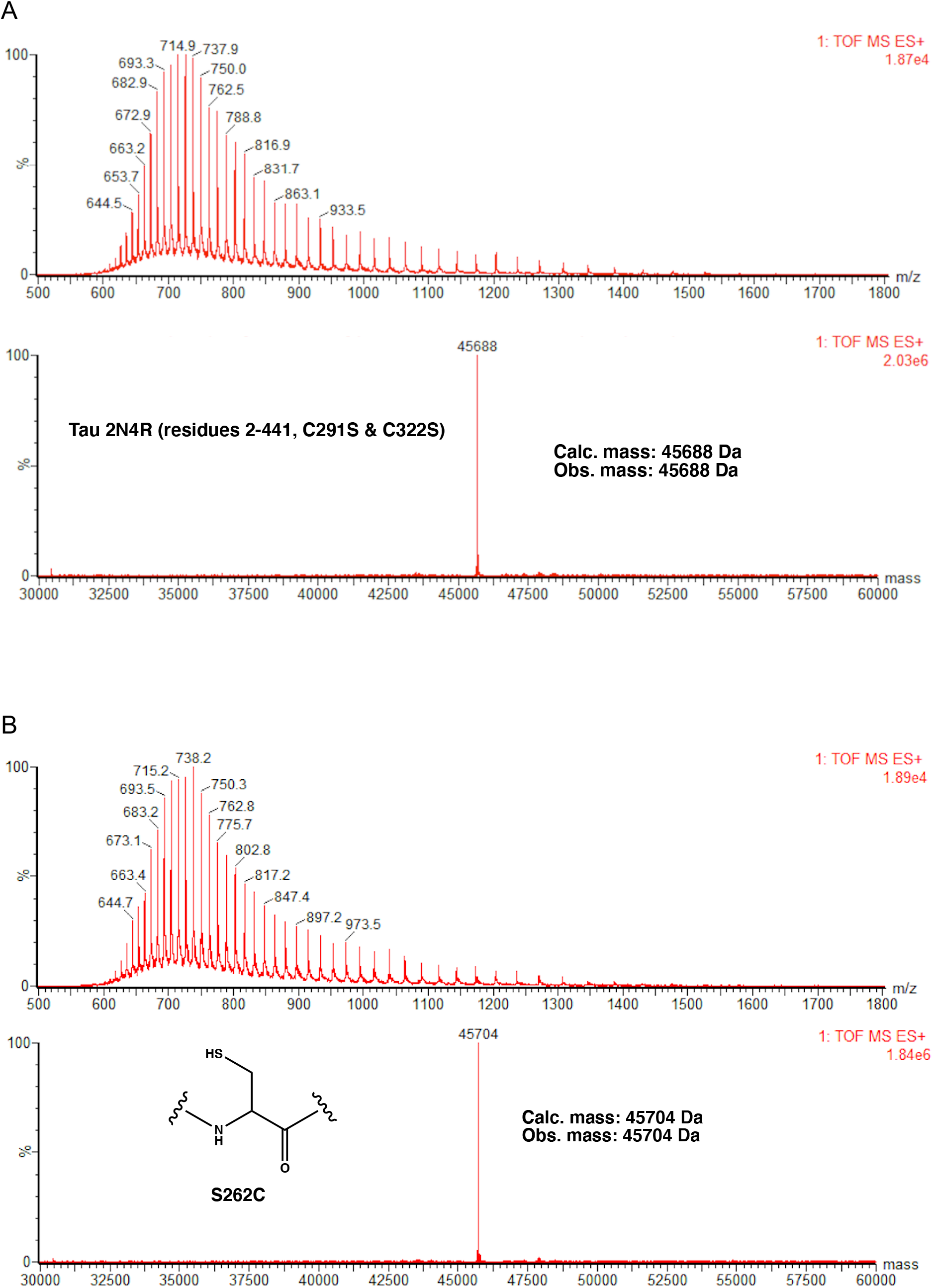
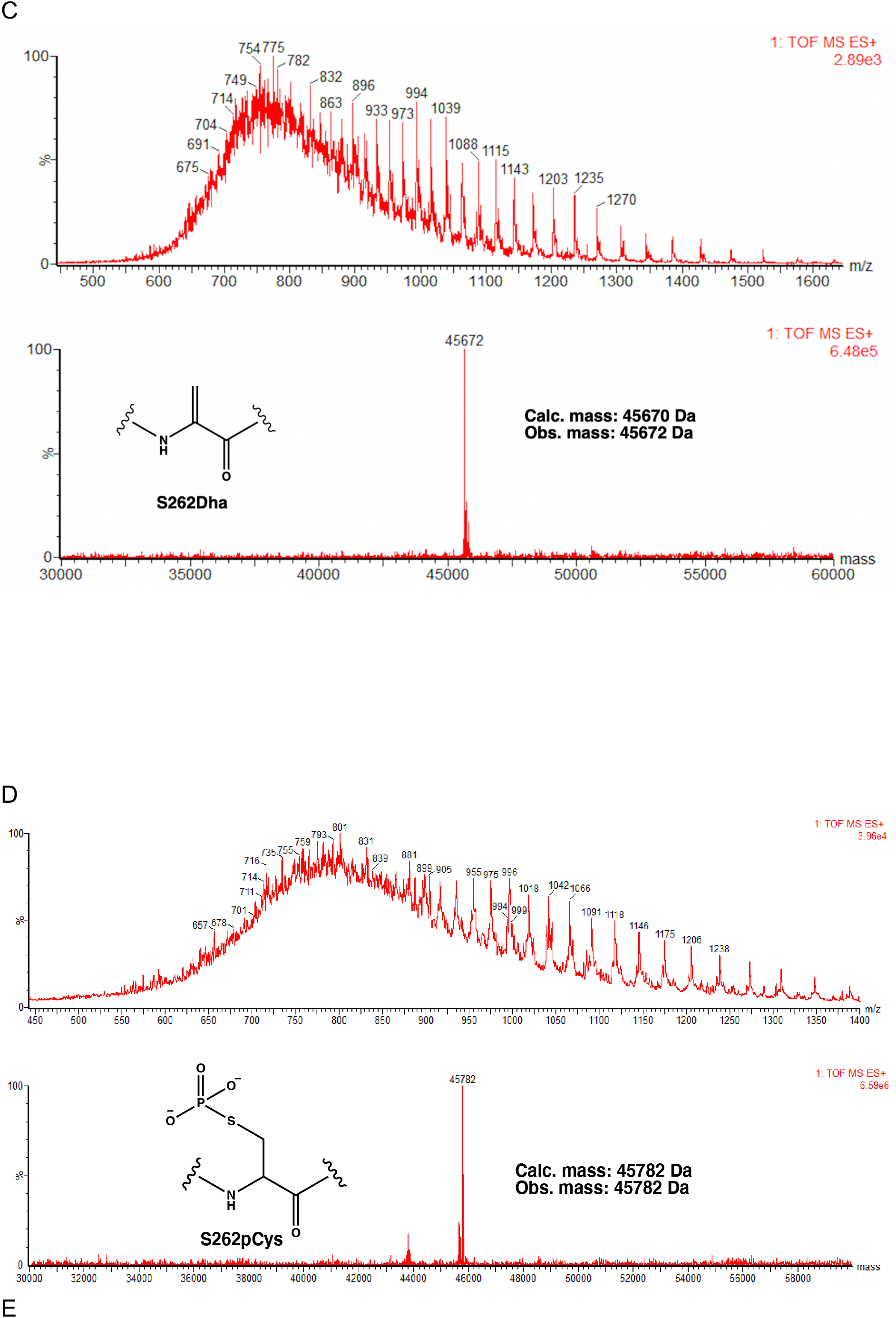
LC-MS spectrum for tau and the phosphorylated chemical mutants and their intermediates. **(A)** LC-MS spectrum for 2N4R tau (residues 2-441, C291S & C322S), all tau variants are residues 2-441. **(B)** LC-MS spectrum for tau(S262C). **(C)** LC-MS spectrum for tau(S262Dha). **(D)** LC-MS spectrum for tau(S262pCys).

**Figure S5.**
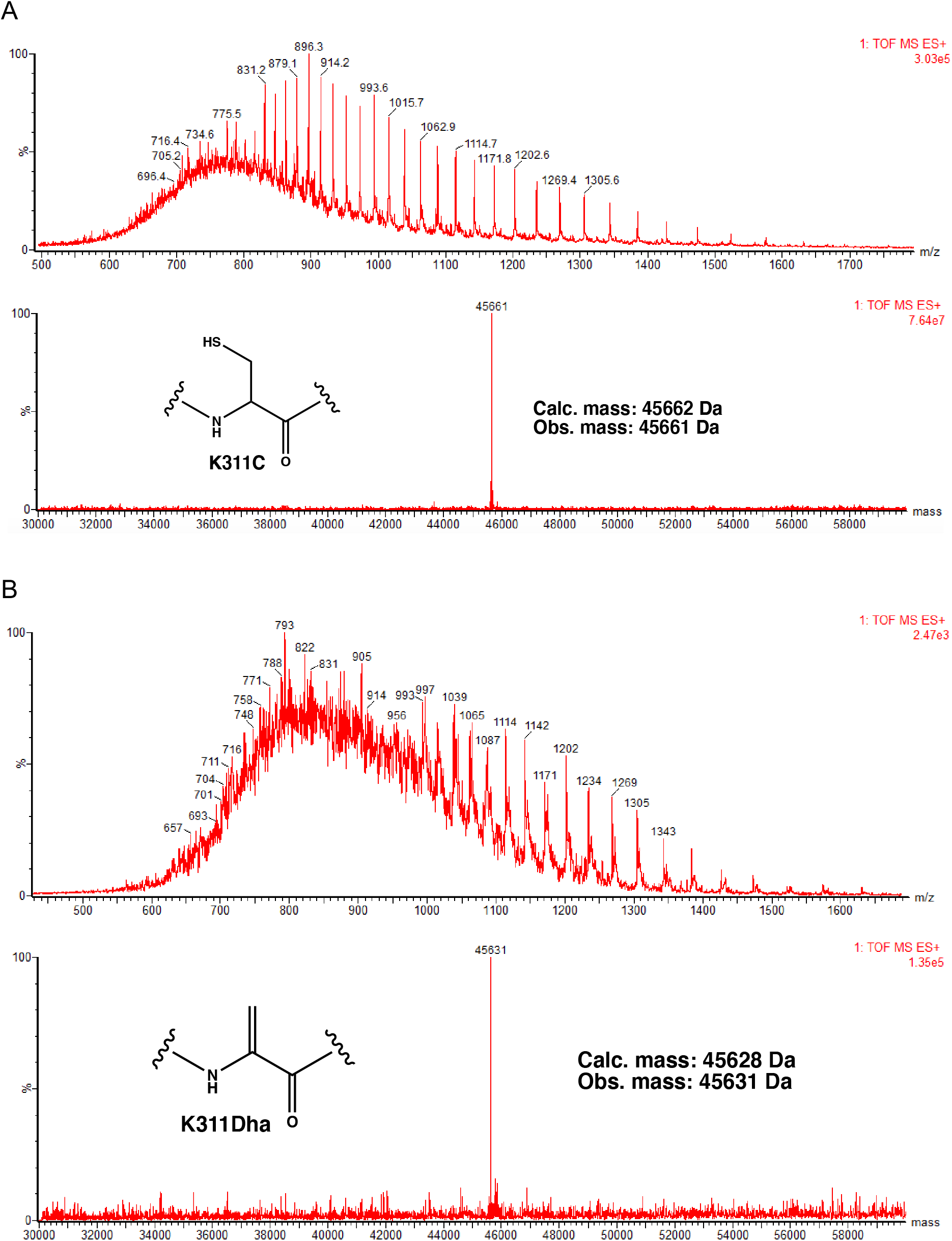
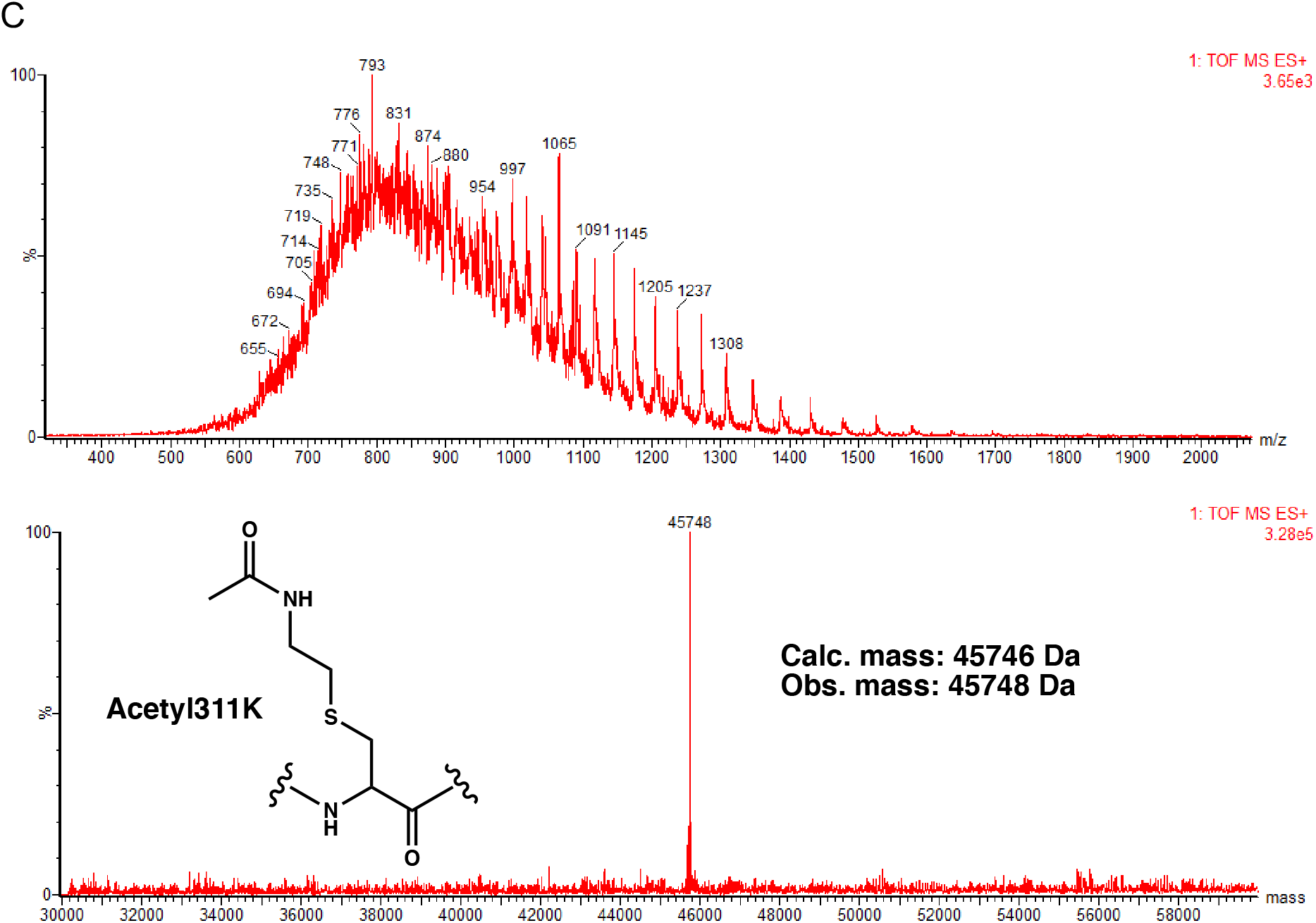
LC-MS spectrum for the acetylated K311 tau chemical mutants and the intermediates. **(A)** LC-MS spectrum for tau(K311C). **(B)** LC-MS spectrum for tau(K311Dha). **(C)** LC-MS spectrum for tau(Acetyl311K).

